# Induction of Chimera States in Hindmarsh-Rose Neurons through Astrocytic Modulation: Implications for Learning Mechanisms

**DOI:** 10.1101/2025.02.10.637588

**Authors:** Fatemeh Azad, Saeed Bagheri Shouraki, Soheila Nazari, Mansun Chan

## Abstract

Chimera states, a form of partial synchronization in neural networks, are characterized by the coexistence of synchronized and asynchronous regions. These states are crucial for various cognitive functions, such as learning and information processing. Conversely, abnormal synchronization—often referred to as hyper-synchronization—can lead to pathological conditions such as epilepsy and Parkinson’s disease. Understanding the mechanisms underlying synchronization can provide valuable insights for developing effective therapeutic strategies for these disorders. Astrocyte, a primary type of glial cell, plays a pivotal role in modulating neural synchrony. They influence the synchronization threshold of neurons by providing feedback through the release of gliotransmitters, promoting group firing of neurons within the astrocyte’s domain. This research aims to explore how astrocytes can facilitate the conversion of hyper-synchronized states into healthy chimera states within neural networks. This process is vital for maintaining normal brain function and may be critical to advancing treatments for neurological conditions. We analyzed how astrocytes can induce chimera states in nonlocally two-dimensional Hindmarsh-Rose neurons, which serve as realistic models of neuronal ensembles. Our findings demonstrate that astrocytes can effectively transition unhealthy hyper-synchronization states into healthy chimera states. Furthermore, by analyzing time spans, spatiotemporal patterns, inter-spike interval distributions (ISI), and phase plane diagrams of 2D H-R neurons, we validated our hypothesis about the crucial role of astrocytes in the development of chimera states. The outcomes may pave the way for innovative therapeutic approaches to restore normal neural activity patterns, ultimately improving patient outcomes in conditions such as epilepsy and Parkinson’s disease.

## 1. Introduction

Recent studies have placed a growing emphasis on synchronization in both natural and artificial complex networks due to the significant impact it has on their functions associated with various behavioural states [1-3]. Numerous neural activities depend on synchronization, which has been detected in several regions of the central nervous system (CNS). Research has demonstrated the critical role that synchronous neuronal play in a variety of brain processes, both in healthy normal conditions (e.g, information processing and cognitive and learning tasks) [4,5], and in unhealthy pathological ones such as Parkinson’s disease due to enhanced synchronization of neurons [6-8]. Phase synchronization [9-12] and generalized synchronization [13-15] are two examples of the several types of synchronization. Phase synchronization refers to the synchronization of the phases of connected systems, without any synchronization of their amplitudes. This is exceptionally relevant for neurological applications, where the precise amplitudes of brain spikes are not as critical as their timing.

In 2002, Kuramoto and Battogtokh [16] identified a specific spatiotemporal pattern that indicates the coexistence of synchrony (coherent behaviour) and asynchrony (incoherence) in a system. This phenomenon, termed the chimera state, has been observed in various scientific systems, including time-continuous chaotic models [17], Van der Pol oscillators [18], FitzHugh-Nagumo neural networks [19], Hindmarsh-Rose oscillators [20], quantum oscillator systems [21], chemical oscillators [22], mechanical oscillators [23], laser systems [24], electronic circuits [25], optoelectronic systems [26], and many others. In addition, chimera states have been shown to exist in neural systems, a finding that raises the possibility of a connection to cognitive disorders, including epilepsy, Parkinson’s disease, and Alzheimer’s disease. Since the aberrant brain rhythms seen in these conditions are mirrored by coexisting coherent and incoherent neuronal activity in chimera states, this knowledge can help develop targeted therapies that seek to restore normal synchronization patterns, potentially reducing symptoms and improving patient outcomes. Furthermore, chimera states are thought to be significant to different activities of the brain because they can describe patterns of partial synchronization. Uni-hemispheric sleep, in which one brain hemisphere is asleep and synchronized while the other is awake and active, has been linked to chimera states in certain animals [27,28]. Recent experiments have demonstrated that the influence of synaptic plasticity, particularly STDP, on chimera states presents another possible use for these states. For example, synaptic plasticity has been shown to break synchronization in a study on multilayer neural networks, resulting in periodic oscillations between layers [29]. Another investigation showed that STDP lengthens the lifetime of Chimera states and expands the range of parameters where they arise. This implies that STDP has a significant impact on neural network dynamics and may impact on learning processes [30]. These applications demonstrate the significant potential of chimera states in neuroscience, ranging from insights into sleep and cognitive dynamics to predictive models for neurological brain disorders.

Recent neurophysiological research has highlighted the critical role of astrocytes, the most prevalent glial cells in the brain, in neural synchrony. Astrocytes modulate the synchronization threshold between coupled neurons, thereby influencing overall neural network dynamics by adjusting neuronal excitability and facilitating forming synchronization and desynchronization of neural networks [31]. They achieve this by releasing gliotransmitters, such as ATP and glutamate, through generating tripartite synapses, which affect synaptic activity [31,32]. By leveraging these regulatory functions, astrocytes can potentially convert hyper-synchronized states, often associated with pathological conditions like epilepsy, into normal chimera states within the same network. In this paper, we suggest that astrocytes play a pivotal role in creating chimera states. This study will investigate this phenomenon and demonstrate that our model supports this concept. Astrocytes control the synchrony of coupled neurons, which help maintain healthy brain function and prevent pathological states. Astrocytes provide feedback to neurons, which can stabilize or destabilize network activity, leading to the emergence of chimera states by creating regions of synchronized and desynchronized activity. Additionally, this research has demonstrated that astrocytes play a crucial role in synaptic plasticity mechanisms, such as spike-timing-dependent plasticity (STDP), by releasing D-serine and modulating the strength and direction of synaptic plasticity. This feedback loop is crucial for balancing long-term potentiation (LTP) and long-term depression (LTD) during learning [33-35]. As a result, astrocytes not only serve as fundamental modulators of neural synchrony but also as vital components in the learning mechanisms that underpin cognitive functions. Thus, in real-world applications, the integration of astrocytes in Spiking Neural Networks (SNNs) leads to low latency, high throughput, and decreased memory consumption, making it highly ideal for resource-constrained settings.

In this research, we address the issue of how astrocytes, acting as the population neural network, might desynchronize nonlocally coupled neurons and generate chimera states. Given the pivotal role of astrocytes in synaptic plasticity and emerging evidence that chimera states are closely linked to learning processes, it is plausible to hypothesize that astrocytes could be active in creating these states. It indicates that neurons do not cease oscillating or begin oscillating in a divergent manner in response to astrocyte stimulation. Actually, the synchrony is broken by the stimulation simply by moving the neurons’ phase by π radians, and with minimal effort, the astrocyte can break the synchrony state and continue to exhibit the appropriate desynchronized behaviour. By adjusting neuronal excitability and facilitating synaptic plasticity, astrocytes may create the conditions necessary for the emergence of chimera states, thereby playing a critical role in the complex interplay between neural synchronization, desynchronization, chimera states and learning. In this paper, we explore the role of astrocytes in inducing chimera states within a neural network of 1000 non-locally coupled two-dimensional Hindmarsh-Rose (2D H-R) neurons arranged on a ring. Firstly, by varying main parameters such as the number of neurons (N), number of nearest neighbours on both sides of each neuron (R), and coupling strength (***σ***), we investigate different behaviour of desynchronization, chimera state, and hyper-synchronization. Then, to investigate the influence of astrocytes, we introduce 500 astrocytes into the network, each receiving input from two randomly selected neurons and forming 500 tripartite synapses by injecting currents into both pre- and post-synaptic neurons. Our findings demonstrate that astrocytes can effectively convert hyper-synchronized states into chimera states by providing feedback to neurons. This feedback mechanism, which can either stabilize or destabilize network activity, leads to the emergence of regions exhibiting both synchronized and desynchronized activities, thereby highlighting the critical role of astrocytes in modulating neural dynamics and promoting complex activity patterns. Ultimately, the validation of all observed states, including asynchronous, hyper-synchronous, and chimera, is accomplished through the analysis of spatiotemporal patterns, membrane potential snapshots of neurons, recurrence plots, and phase-plane diagrams. Additionally, mean inter-spike interval (ISI) and global order parameters are measured to confirm these states. The significance of our findings lies in the potential applications for understanding complex brain functions and disorders characterized by abnormal synchronization. Our work provides theoretical support for future studies on the coexistence of multiple states in the brain and paves the way for future research into therapeutic strategies targeting astrocytic functions to restore normal brain activity patterns by clarifying the mechanisms through which astrocytes modulate neural activity.

By leveraging the relationship between chimera states and synaptic plasticity mechanisms including LTP and LTD, the proposed SNN shows how chimera states can function as a basic building block for learning. By integrating these dynamics, the SNN highlights how the presence of coherent and incoherent neural activities can facilitate adaptive learning processes. This paper’s structure is set up as follows: In Section 2, the 2D H-R model of neuron, astrocyte models and their phase portraits are presented mathematically. In Section 3, the construction of the whole proposed neural network of the study considering the presence of astrocytes is discussed. Section IV discusses the findings in detail as well as the function of astrocytes in fostering spike-timing-dependent plasticity (STDP) and the chimera state. Section V provides a summary of the findings and recommendations for further investigation.

## 2. Models and Methods

In this section, to investigate the collective dynamics of 2D H-R neurons and astrocytes, mathematical models are given. Furthermore, phase portraits are utilized to visually depict the system’s dynamics, providing critical insights into the stability and interaction patterns of neurons and astrocytes under various initial conditions.

### 2.1. 2D-HR Neuron Model

Here, two first-order ordinary differential equations (ODEs) describe the two-dimensional Hindmarsh-Rose (H-R) neurons. Under periodic boundary conditions, these equations describe the dynamics between the membrane potential and a related variable pertaining to ionic currents across the membrane [36]. We examine a network of (N) nonlocally coupled Hindmarsh-Rose neurons that are connected to their (R) nearest neighbours on both sides. The equation governing this system is as follows:

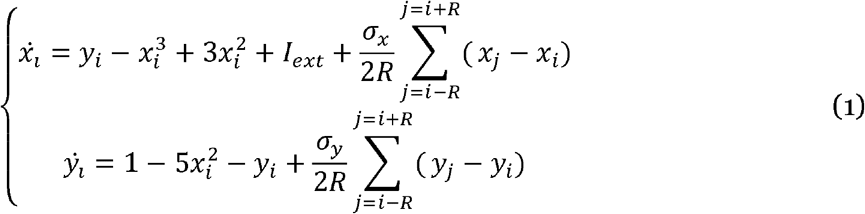

Where *x*_*i*_ represents the membrane potential of the *i*th HR neuron, *y*_*i*_ corresponds to various physical quantities related to the conductance of ion currents across the membrane, and *I* _*ext*_ is the external stimulus current, which is set to zero as the coupling between the neurons serves as the stimulus current. Each neuron has coupling strengths *σ*_*x*_ and *σ*_*y*_ with its *R* nearest neighbours on both sides. This setup introduces nonlocality in a ring network topology, facilitated by periodic boundary conditions for the ODE system. Fig. 1(a) illustrates the 2D phase portraits of the 2DH-R neuron model for *I* _*ext*_ =0. The initial condition for *x* and *y* is selected through random numbers on a unit circle (*x*^2^+*y*^2^ = 1).

**Figure 1:**
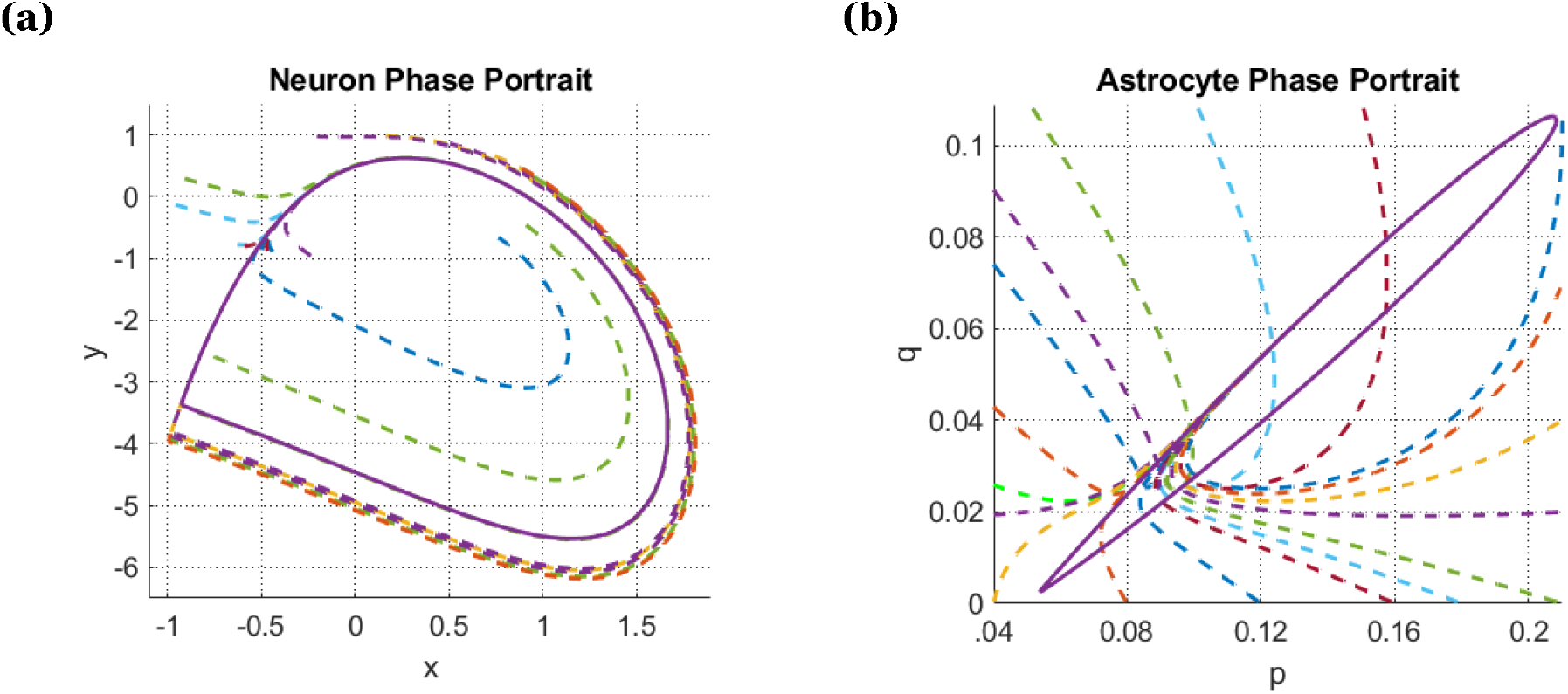
The 2D phase planes of the neuron and astrocyte model, based on Eq. (1) and Eq. (2). **(a**) 2D phase portraits of the 2D H-R neuron model for by changing initial conditions randomly. **(b)** The phase plane of astrocyte states in (p − q) space by changing initial conditions randomly.

### 2.2. Astrocyte Model

An astrocyte-inspired stimulator [37] has been used to effectively desynchronize coupled 2D H-R neurons, demonstrating a significant advancement in neuromorphic engineering. This bioinspired astrocyte stimulator operates without inducing adverse effects on the intrinsic behaviours of the neurons, such as halting, annihilating, or instigating divergent oscillations, which are critical for maintaining neuronal functionality and stability [37, 38]. By leveraging the principles of spiking neural networks (SNNs), which emulate the processing capabilities of biological neurons, the astrocyte-inspired stimulator improves the control over neuronal dynamics, allowing for precise modulation of oscillatory behaviour in coupled neuronal networks (Qiu et al., 2018). This innovative approach not only underscores the potential of bioinspired designs in neuromorphic applications but also opens avenues for further exploration in controlling complex neural interactions without compromising the crucial characteristics of neuronal activity. Based on the original model of astrocyte, the structure of the bioinspired controller [37] is as follows:

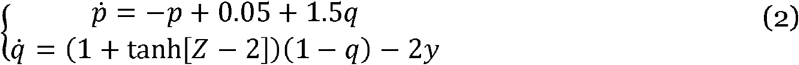

in which *p* and *q* are the astrocyte states. Based on the observation Z (input of astrocyte stimulator) of the network states, astrocyte produces the appropriate stimulation. Thus, Z acts as an astrocyte input, accumulating the neuronal membrane potential via system (3).

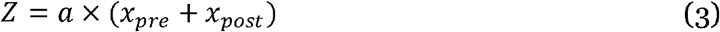

Where *x*_*pre*_ and present *x*_*post*_ the membrane voltages of pre- and post-synaptic neurons, respectively, which are connected to an astrocyte through a tripartite synapse. The parameter a denotes the astrocyte input gain, which is used to adjust *Z* to maintain the hyperbolic tangent function within its linear region. To show the astrocyte states’ phase portrait of in (p-q) space (Fig. 1(b)), Z, the astrocyte input, is configured as *Z* =sin (0.1*t*).

## 3. Network Topology

In this paper, we construct a cortical neuronal network, based on tripartite synapse using N=1000 nonlocally coupled identical H-R neurons and M=500 identical astrocyte-inspired stimulator. Each astrocyte receives input from pre- and post-synaptic neurons, based on Eq. (3) and then produce two currents *I*_1_ and *I*_2_ to apply to the same neurons as follows:

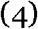

Where and serve as feedback gain produced by the astrocyte, and p is the output of the stimulator. The close loop of a single tripartite synapse, which is made up of two neurons and one astrocyte is exposed in Fig. 2 (dashed square box). This information is used to characterize the close-loop system as follows:

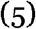

**Figure 2:**
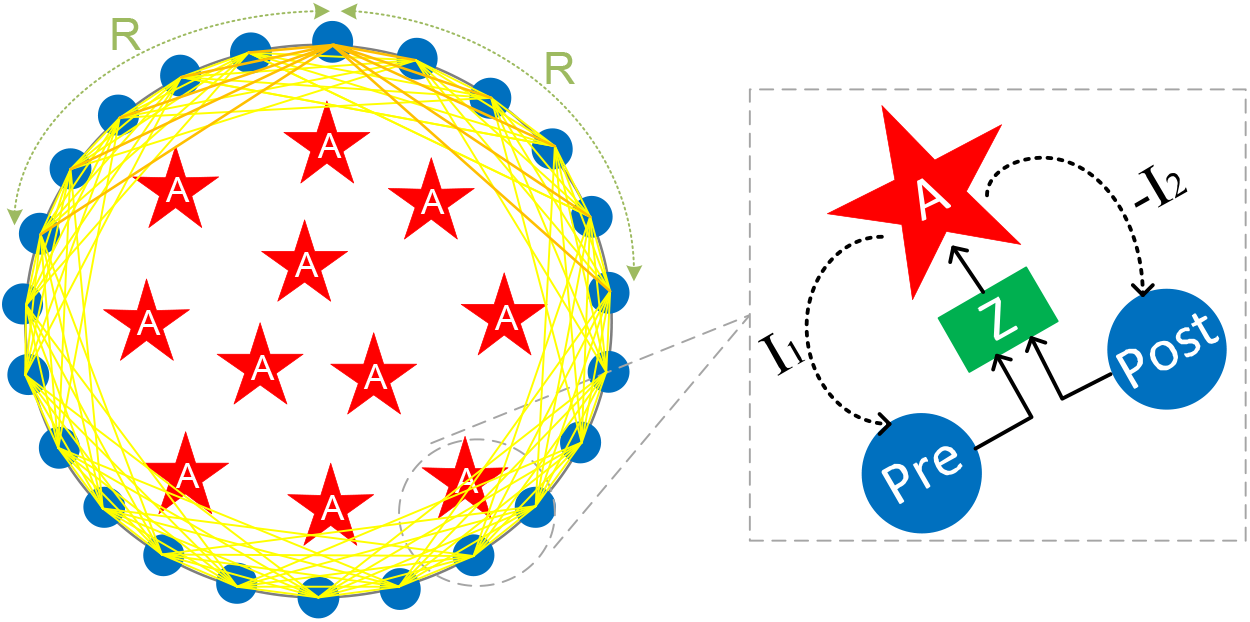
The Schematic view of the whole network and a depiction of how astrocyte modulate synaptic currents highlighting the primary routes for interactions between neurons and astrocytes.

Finally, the two coupled neurons are controlled (desynchronized) by p via the parameters and, which are also known as stimulation. Among the gliotransmitters released by astrocytes, glutamate has an excitatory effect while ATP has an inhibitory effect [37]. A positive sign for excitation and a negative sign for inhibition are used to model this reality. In the rest of this paper, we applied 500 tripartite synapses to build the whole network dynamically. Figure (2) illustrates the whole network topology including N neurons arranged on a ring with R nearest neighbours on both sides of each neuron. The order of influences and interactions within a tripartite synapse is depicted in this diagram: when information from the neural network reaches the presynaptic terminal, the process starts. Subsequently, neurotransmitters are released and spread across the synaptic cleft binding to receptors on the astrocyte membrane. This binding causes the astrocyte to release gliotransmitters, which in turn alter the likelihood that neurotransmitters will be released at both the pre- and post-synaptic terminals.

## 4. Numerical Results and Discussion

The selection of values for coupling strength(***σ***_***x***_ and ***σ***_***y***_), range of coupling (*R*), and the neural network size (*N*) is crucial for observing different chimera states. Here is an explanation of how these parameters influence forming chimera states: When the coupling strength is low, the influence of neighboring neurons on each neuron is weak. This can lead to more localized and less synchronized behaviour, enhancing forming incoherent domains. When the coupling strength is high, the influence of neighboring neurons is more substantial, promoting synchronization across the network. This can lead to more coherent behaviour and fewer incoherent domains. R determines the numbers of nearest neighbours on either side that each neuron interacts with. A more extensive range of coupling means each neuron interacts with more neighbours. This tends to promote global synchronization, potentially reducing the numbers of incoherent domains. A smaller range of coupling means each neuron interacts with fewer neighbours. This can lead to more localized interactions and enhance producing multiple incoherent domains. With a smaller system size (N), the complexity of the network is limited, and the patterns of coherence and incoherence are simpler. Increasing the system size introduces more complexity, allowing for generating multiple incoherent domains and more intricate chimera states.

In this paper, the Euler method is employed to numerically simulate the dynamical equations at a time step of *dt* = 0.01. The initial conditions of the variables are fixed and randomly dispersed on the unit circle(***x***^2^+***y***^2^=1) in all simulations. First, to illustrate the different behaviours of asynchronization, chimera states, and complete synchronization, we consider the astrocyte-free neural network on a one-dimensional ring, in which every neuron is connected to its 2*R* nearest neighbours for different values of coupling strength and neighboring range (*R*) (Fig. 3-6). In Fig. 3 (a), the blue and red colour in time series panel shows the activity of neurons numbers 200 and 300, the green one illustrates the activity of neuron number 400. For 1000 neurons with R=350, and ***σ***_***x***_=***σ***_***y***_=0.73, the network forms multiple incoherent regions, known as multiheaded chimera state [39]. When we increase the coupling strength t0 ***σ***_*x*_ ***σ***_***y***_=0.15, neurons will be synchronized (Fig. 3 (b)). Research indicates that as the coupling range decreases, the complexity of the chimera states can increase, allowing for multiple coherent and incoherent domains to coexist. Multiple regions of incoherence and coherence arise as chimera states with a decreased coupling range (R) in Fig. 4(a). Fig. 3(b) and Fig. 4(b) illustrate cases of hyper-synchronization patterns within neural network. Hyper-synchronization occurs when neurons within a network fire in a highly coordinated manner, leading to excessive synchronous activity. This phenomenon is often observed in various neurological disorders, such as epilepsy, where it manifests as seizures [40-42]. The state of hyper-synchronization is primarily influenced by the coupling strengths between neurons. When the coupling strengths are too high, the interactions between neurons become excessively strong, causing them to fire in unison. This high degree of synchronization can disrupt normal brain function, leading to pathological conditions. In the context of our study, the coupling strength refers to the synaptic connections and the degree of influence one neuron exerts on another. As the coupling strength increases, the likelihood of neurons firing together also increases, resulting in the hyper-synchronized state depicted in Figures 3(b) and 4(b). This state is characterized by losing the normal, dynamic balance of neuronal activity, which is critical for healthy brain function. Understanding the mechanisms underlying hyper-synchronization is crucial for developing therapeutic strategies to mitigate its effects.

**Figure 3:**
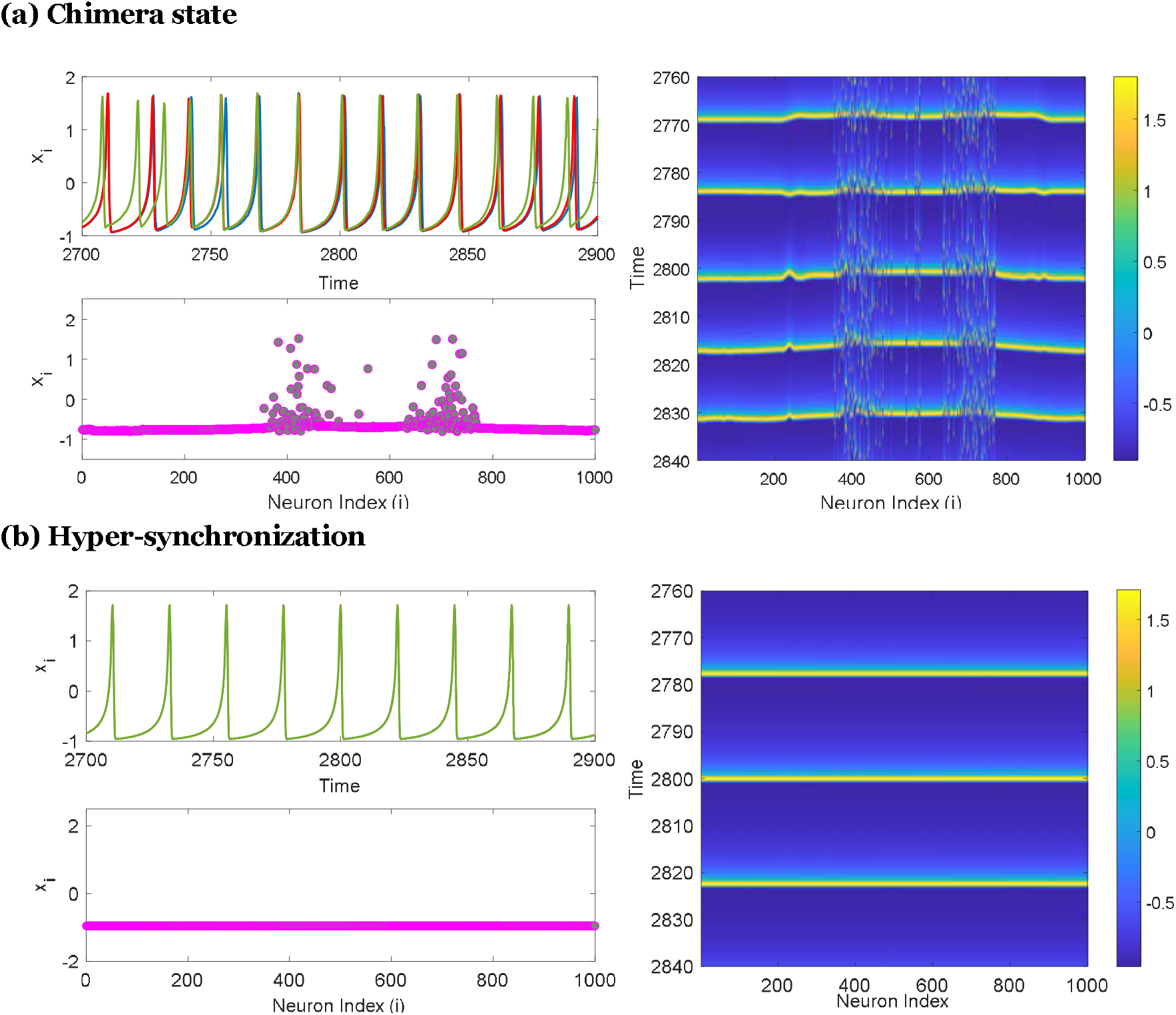
Temporal sequences, snapshots, and spatiotemporal patterns of the neural network without astrocyte (Eq. 1) under varying coupling strength and 0 for *I* _*ext*_ = 0 N=1000, R=350. **(a)** Chimera state at 2780 sec (snapshot) with *σ*_*x*_ = *σ*_*y*_ = 0.073, time series of neuron 200 (blue) and 300 (red) showing the same phase but neuron number 400 (green) has different phase. **(b)** Hyper-synchronization state at 2780 sec (snapshot) with *σ*_*x*_ = *σ*_*y*_ = 0.15, time series of neuron 200 (blue), 300 (red), and number 400 (green) illustrating the same frequency.

**Figure 4:**
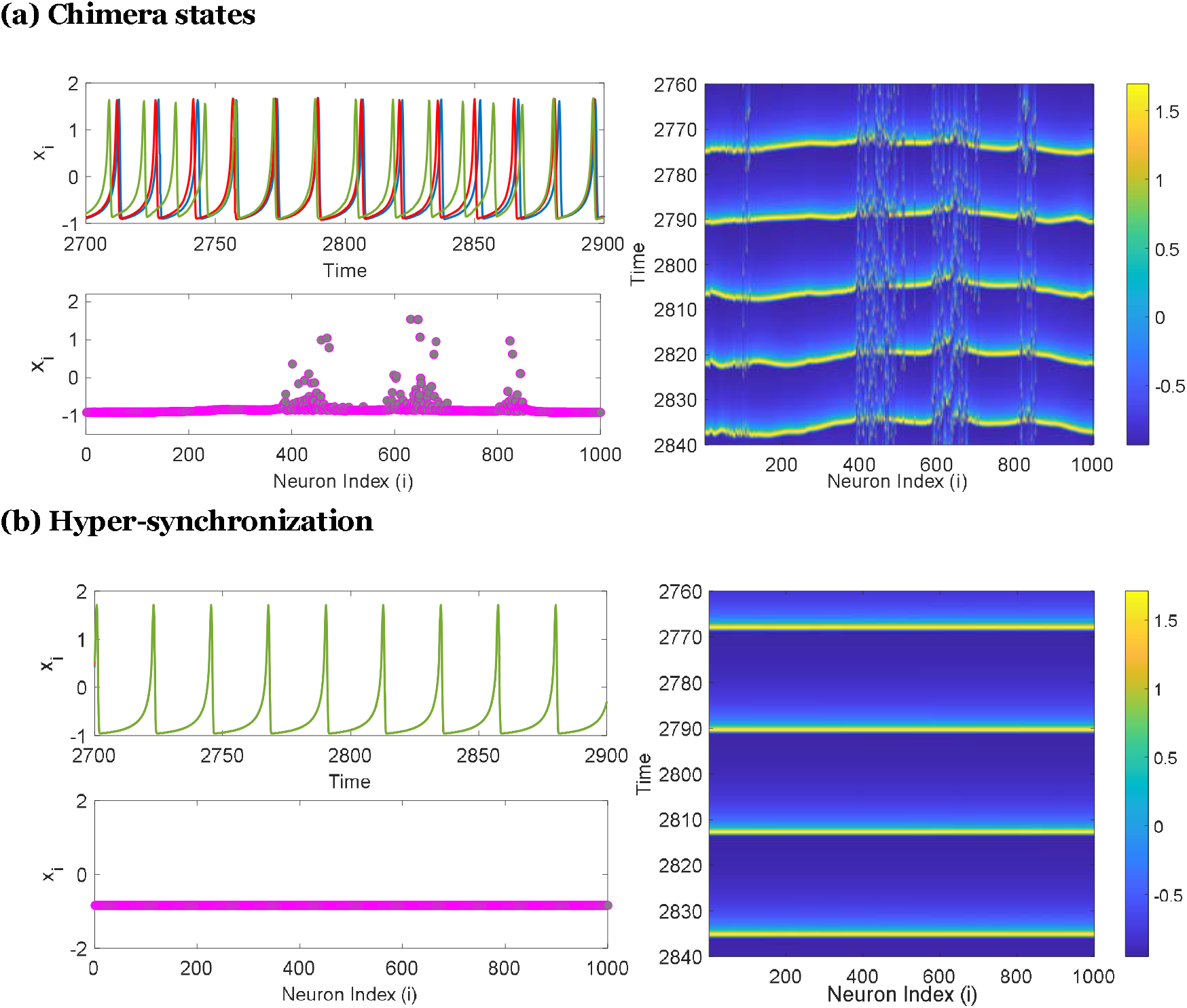
Temporal sequences, snapshots, and spatiotemporal patterns of the neural network without astrocyte (Eq. 1) under varying coupling strength and *I* _*ext*_ = 0 for N=1000, R=250. **(a)** Chimera state at 2780 sec (snapshot) with ***σ***_*x*_=***σ***_*y*_=0.052, time series of neuron 200 (blue) and 300 (red) showing the same phase but neuron number 400 (green) has different phase. **(b)** Hyper-synchronization state at 2780 sec (snapshot) with ***σ***_*x*_=***σ***_*y*_= 0.122, time series of neuron 200 (blue), 300 (red), and number 400 (green) illustrating the same frequency.

By modulating the coupling strengths, either through pharmacological interventions or neuromodulation techniques, it may be possible to restore normal neuronal activity and alleviate symptoms associated with hyper-synchronization.

In the context of epilepsy and other brain disorders characterized by hyper-synchronization, adding astrocytes can influence neuronal network dynamics. For instance, adding astrocytes to a hyper-synchronized network can help restore balance by modulating neurotransmitter levels and buffering extracellular potassium [43]. This intervention can lead to the emergence of chimera states, where some regions of the system exhibit synchronized activity while others remain desynchronized. These chimera states can reduce the overall pathological synchronization, potentially alleviating symptoms associated with hyper-synchronization in brain disorders. In this way, we considered a case by adding 500 astrocyte-inspired stimulators to illustrate the effect of their crosstalk with neurons in converting hyper-synchronization to multiple chimera states.

Fig. 5 illustrates how 500 tripartite synapses formed by 500 astrocytes can convert hyper-synchronization behaviour of spiking neural network to chimera states. Desynchronization as the primary control goal is not achievable without choosing an appropriate set of controller parameters, particularly the values of *λ*_*1*_ and *λ*_*2*_ is selected. In Fig. 5, the snapshots of 1000 neurons at 3933 sec, along with the spatiotemporal patterns of neurons in the presence of astrocytes during the interval from 3900-4000sec, illustrate the crucial role of astrocytes in inducing chimera states. This occurs in a network with the same coupling strength as that of hyper-synchronization states, indicating that the values of *σ*_*x*_ and *σ*_*y*_ are the same as the values in hyper-synchronization states (*σ*_*x*_ = *σ*_*y*_= 0.122 when R=250 and*σ*_*x*_ = *σ*_*y*_ = 0.15 when R=350). Furthermore, the numerical results given in this part are examined using the recurrence quantification analysis. The network’s spatiotemporal behaviour, which traverses the same area of the phase space, is virtualized in the recurrent plots. One can compute the recurrent analysis as follows:

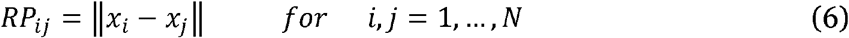

where *N* is the number of neurons and ‖. ‖ is the Euclidean distance and this parameter, ***RP*** _***ij***_ has been normalized. The recurrent plots for various coupling strength values (σ) are displayed in Fig. 6.

**Figure 5:**
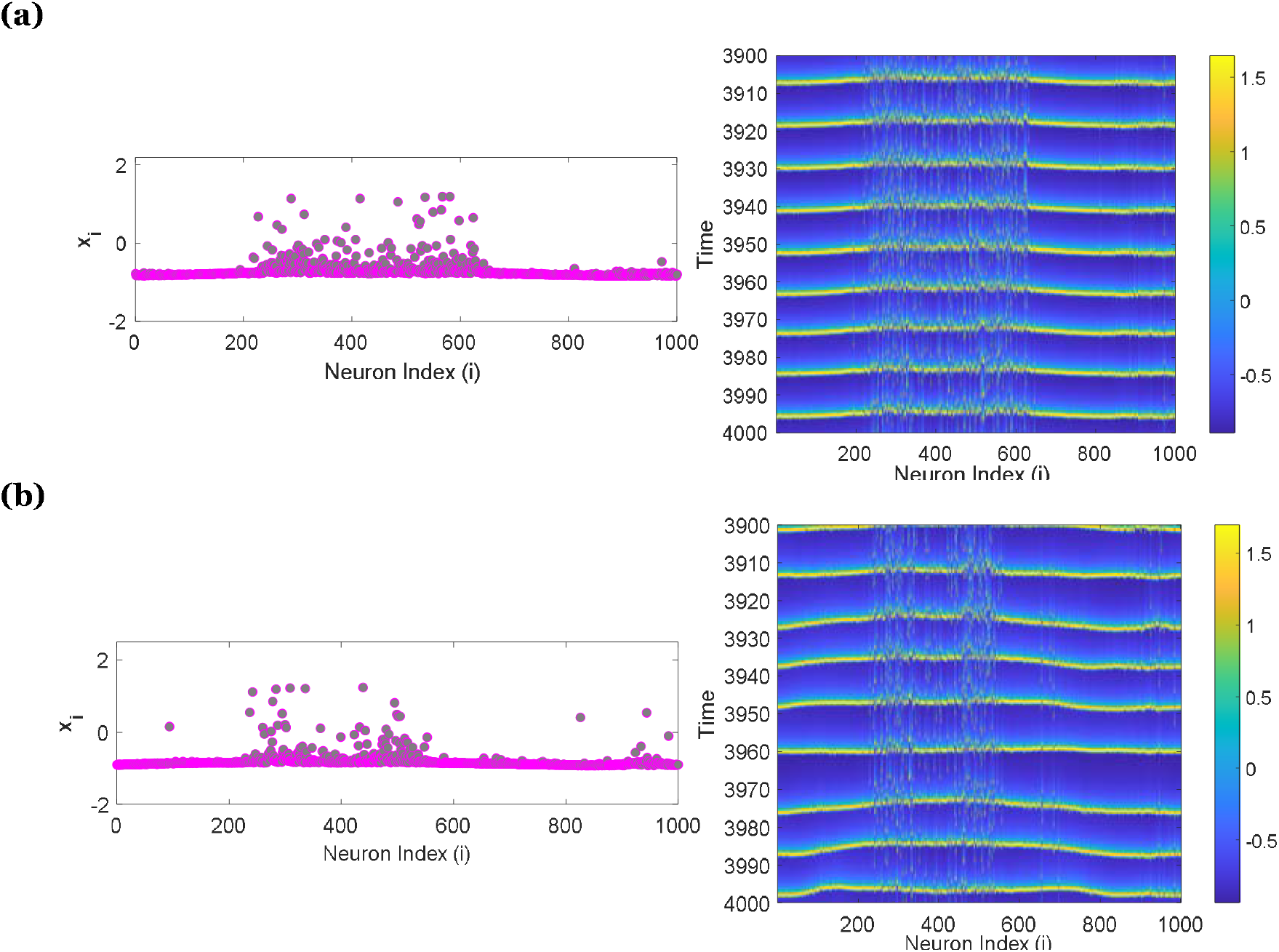
The snapshots and spatiotemporal patterns of the neurons in the neural network with considering astrocytes (Eq. 5) for *λ*_1_ = 0.05 and *λ*_2_ = 0.15 under varying coupling strength and *I* _ext_ = 0. **(a)** Chimera state for N=1000, R=350, σ_*x*_ = σ_*y*_ = 0.15. **(b)** Chimera state for N=1000, R=250, *σ*_*x*_ = *σ*_*y*_ = 0.122.

**Figure 6:**
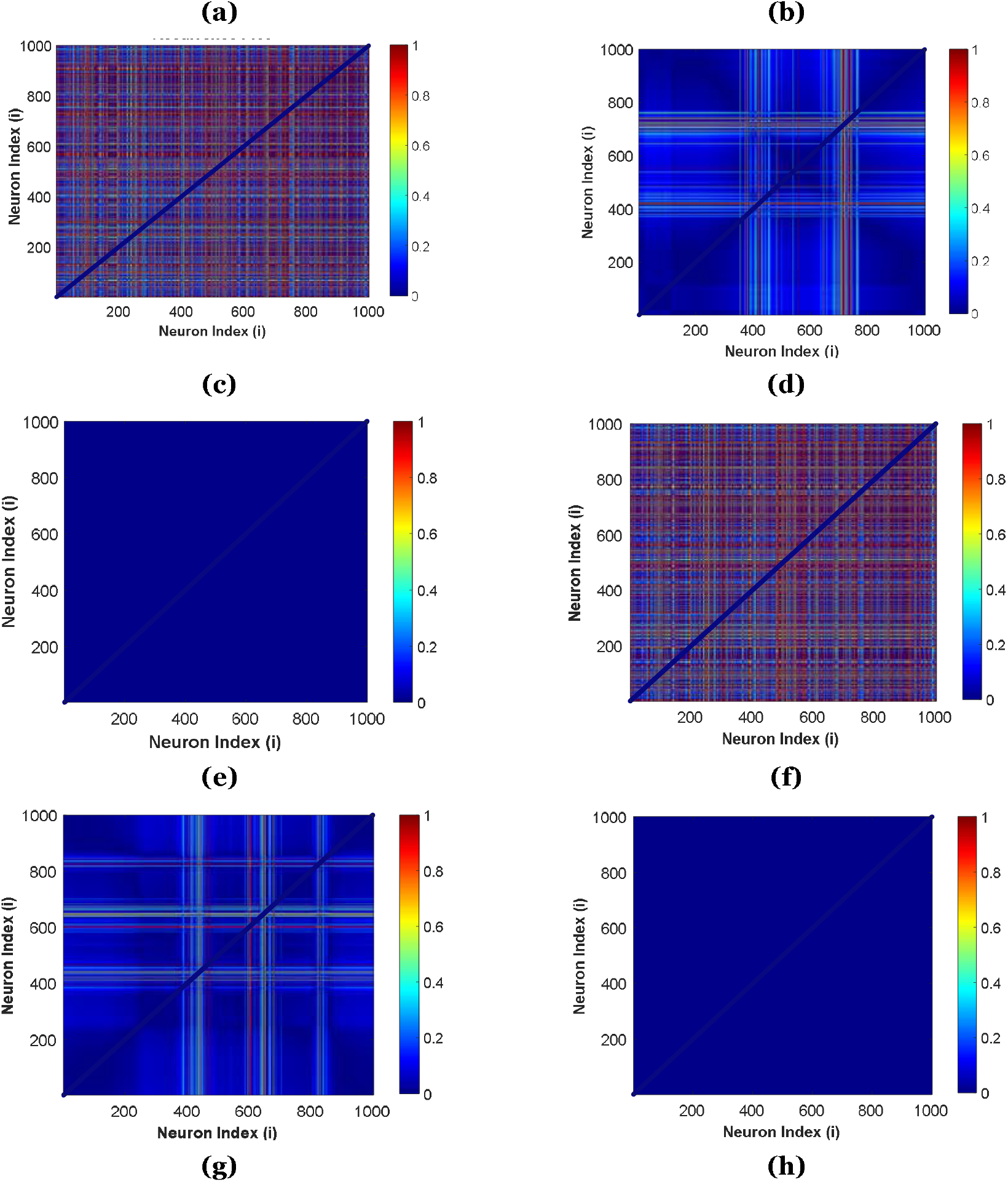
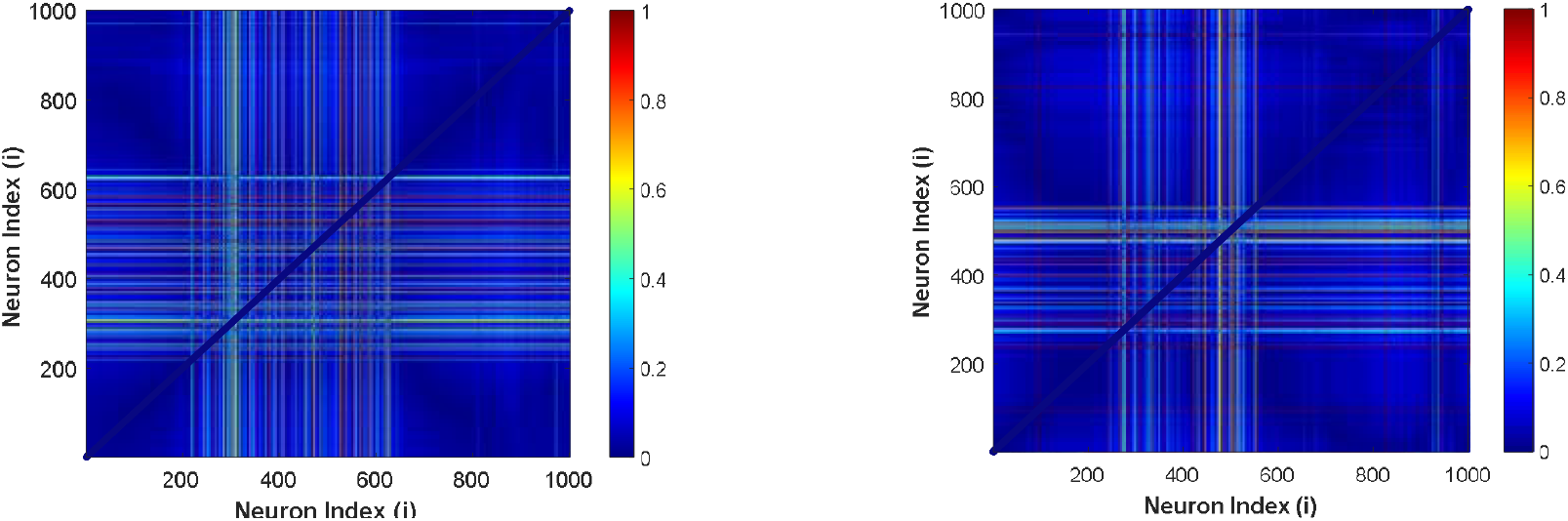
Recurrence plot. (a) N=1000, R=350, (b) N=1000, R=350, ***σ***_*x*_ = ***σ***_*y*_ = **0.073**.(c) N=1000, R=350, ***σ***_*x*_ = ***σ***_*y*_= **0.15**. (d) N=1000, R=250, ***σ***_*x*_ = ***σ***_*y*_= **0.00001**. (e) N=1000, R=250, ***σ***_*x*_ = ***σ***_*y*_= **0.052**. (f) N=1000, R=250, ***σ***_*x*_ = ***σ***_*y*_= **0.122**. (g) N=1000, R=350, ***σ***_*x*_ = ***σ***_*y*_= **0.15**. with astrocytes. (h) N=1000, R=250, ***σ***_*x*_ = ***σ***_*y*_= **0.122**. with astrocytes.

For larger values of the coupling strength (*σ*), hyper-synchronization in the network arises by varying coupling strength (*σ*). Fig. 6 (a) and (d) illustrates the incoherent behaviour of the network without any structures when the coupling strength is adjusted to *σ* = 0.00001. Structures in the recurrent plots of Figures 6(b), (e), (g), and (h) show introducing chimera states with simultaneous synchronous and asynchronous network behaviour. The colour box indicates that the neurons with asynchronous behaviour are represented by red regions, and the synchronized neurons are represented by blue sections. The complete synchronous behaviour (hyper-synchronization) of the network is shown in the recurrent plot of Figures 6(c) and (f).

To compare how the presence of astrocytes influences the synchronization dynamics of the neural network, the global-order parameter against the coupling strength (*σ*_*x*_ = *σ*_*y*_ = *σ*) has been illustrated in Fig. 7(a) and (b). The spatiotemporal average of the local-order parameters is known as the global-order parameter . This parameter (ℛ) functions as a synchronization index, providing a quantitative assessment of how synchronized the neural network is:

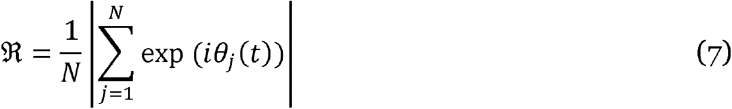

where *θ*_*j*_ (*t*) is the phase of *j*th neuron, calculated by:

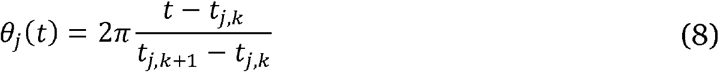

where *t*_*j,k*_ is the time of kth spike of jth oscillator or neuron and *j*=1,2,…,*N*. The value of (ℜ) equals 1 when the network reaches complete synchronization. Therefore ℜ < 1 illustrates that the system is not completely synchronized. In this experiment, we plot the global-order parameter (ℜ) against the coupling strength (σ) to compare two distinct states with different *R* (number of nearest neighbours for each neuron). This comparison allows us to observe how the presence of astrocytes influences the synchronization dynamics of the neuronal network. Averaging 16 trials with randomly chosen initial condition sets and a 4000-millisecond run-time yields each panel in Figure 7. There is an increase in the coupling strength (σ) within the range of 0 ≤ σ ≤ 1.5. In the context of network dynamics, the observed upward trend in the global order parameter suggests that synchronization within the network is positively influenced by an increase in coupling strength. Specifically, it appears that the network is capable of achieving complete synchronization at elevated coupling strengths. Notably, as illustrated in Fig. 7(a) and (b), the global order parameter exhibits a local minimum within the range of 0.04 ≤ σ ≤ 0.1. This phenomenon is attributed to the influence of astrocytes, which alter the range and effectively expand it. Within this interval, an intriguing deviation from the overall trend is observed: the degree of synchronization diminishes as the coupling strength is augmented. This local minimum may serve as a potential indicator of the emergence of a chimera state within the network. Such behaviour has also been documented in heterogeneous Kuramoto network models characterized by both attractive and repulsive coupling interactions [16].

**Figure 7:**
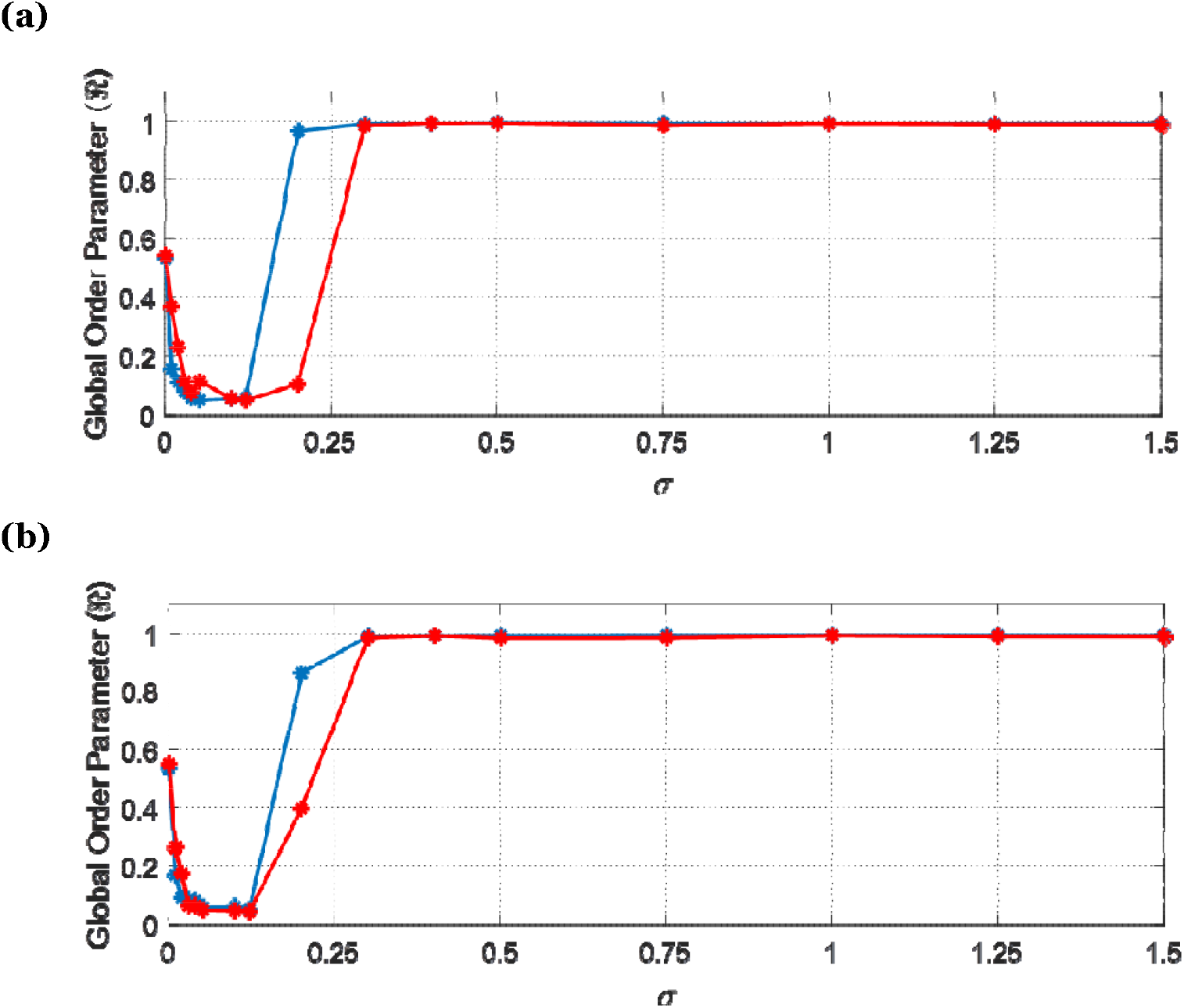
Variation of the global-order parameter () with changing coupling strength (σ) for networks with (red) and without (blue) astrocytes, with N=1000 and M (number of astrocytes=500). **(a)** *R*=250. **(b)** *R*=350. Synchronization is achieved at more decisive couplings while in the presence of astrocytes, the network reaches complete synchronization at higher σ.

In Fig. 7, two distinct scenarios are compared. In panel (a), the synchronization dynamics of neurons are analyzed with (red) and without (blue) astrocytes when R = 250. Panel (b) further examines the network at R = 350. The results reveal that, without astrocytes, the network achieves synchronization at lower coupling strengths (σ). In contrast, when astrocytes are present, the network necessitates higher coupling strengths to attain complete synchronization. Through the analysis of these plots, we gain valuable insights into the role of astrocytes in modulating network dynamics and neuronal synchronization. It appears that astrocytes increase the threshold for synchronization, indicating their regulatory function in neuronal network behaviour. This finding is significant as it underscores the complex interplay between neurons and astrocytes, which may have important implications for understanding brain function and developing treatments for neurological disorders.

The interaction between neurons and astrocytes has garnered significant attention in neuroscience, particularly regarding how astrocytic activity influences neuronal dynamics and interspike interval (ISI) distributions. The introducing astrocytes into a network of nonlocally coupled neurons is expected to alter the mean ISI due to the complex signaling mechanisms that astrocytes employ, primarily through the release of neurotransmitters like glutamate and the modulation of calcium signaling. Evidence from Silchenko and Tass [44] suggests that random calcium oscillations in astrocytes can cause simultaneous bursting activity in neurons by causing synchronous responses in nearby neurons. This phenomenon implies that the presence of astrocytes can lead to coherent oscillations among neurons, potentially affecting the distribution of ISIs. Furthermore, Wallach et al. [45] discuss how astrocytic glutamate release can filter neuronal activity, which may also contribute to changes in ISI distributions by modulating the excitability of neurons. Moreover, the work of Oku et al. [46] highlights that rhythmic calcium fluctuations in astrocytes can precede neuronal activity, indicating a temporal relationship that could influence ISI patterns. This rhythmicity suggests that astrocytes might play a role in synchronizing neuronal firing, which could lead to a more regular ISI distribution. In Fig. 8 and 9 (first rows), when the coupling strength (σ) is 0, every neuron in the network is either in a resting or spiking state because of various initial conditions.

**Figure 8:**
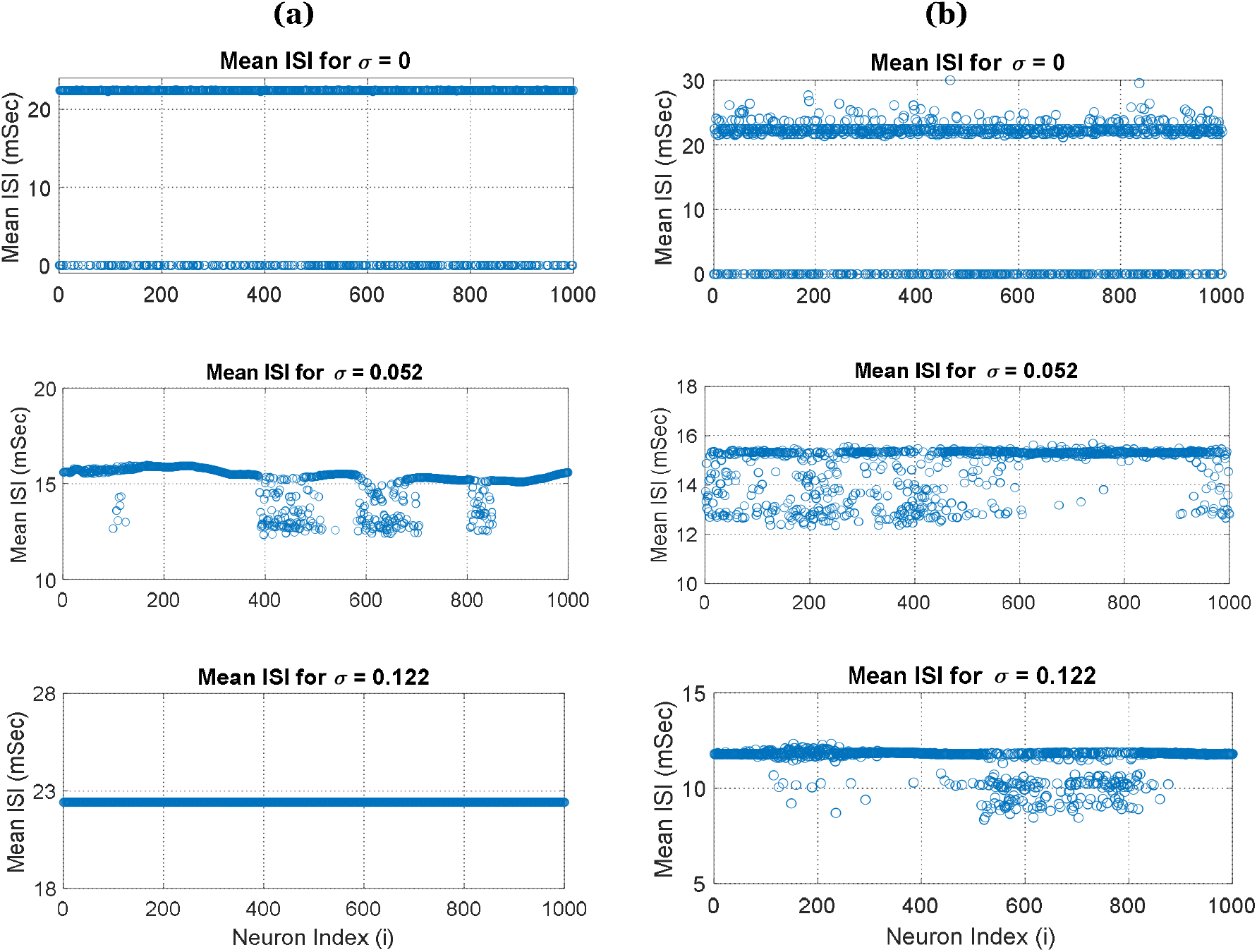
The network’s mean ISI distributions for 1000 neurons of the network, comparing scenarios with and without the presence of astrocytes while varying the coupling strength (σ). **(a)** Mean ISI for different coupling strength (σ = 0, 0.073, and 0.15) without astrocytes. **(b)** Mean ISI for the same coupling strength (σ = 0, 0.073, and 0.15) when astrocytes included in the system. Here, the network demonstrates a range of synchronization states, including asynchronization, chimera state, and completely synchronization, corresponding to the specified σ without presence of astrocytes in the network. When astrocytes added to the network, for more extensive σ, some neurons behave differently. Thus, astrocyte convert hyper synchronization state to chimera (Third row (b)). Here, the value of *N* = 1000 and *R* is 250.

**Figure 9:**
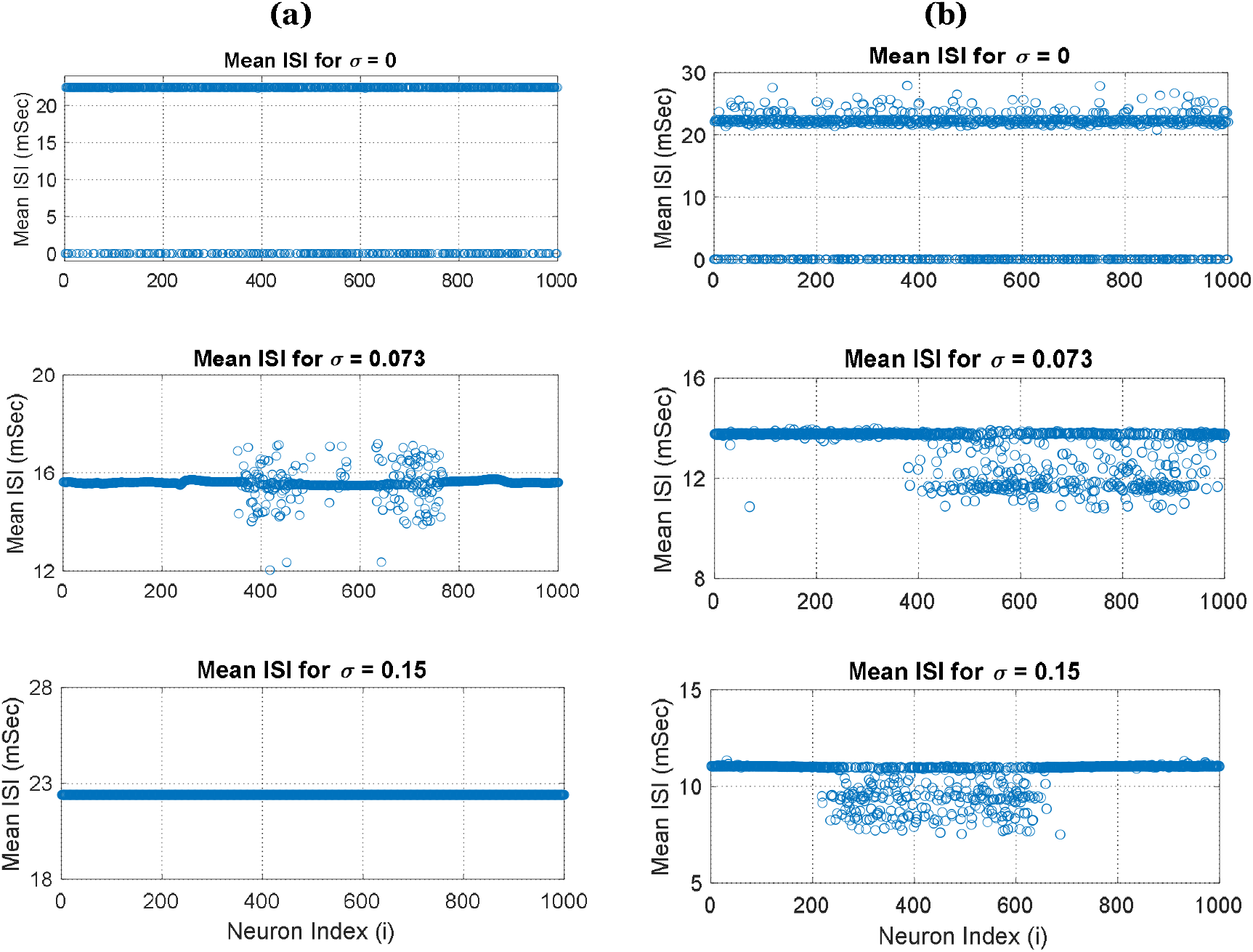
The network’s mean ISI distributions for 1000 neurons of the network, comparing scenarios with and without the presence of astrocytes while varying the coupling strength (σ). **(a)** Mean ISI for different coupling strengths (σ = 0, 0.073, and 0.15) without astrocytes. **(b)** Mean ISI for the same coupling strengths (σ = 0, 0.073, and 0.15) when astrocytes included in the system. Here, the network demonstrates a range of synchronization states, including asynchronization, chimera state, and completely synchronization, corresponding to the specified without presence of astrocytes in the network. When astrocytes added to the network, for larger, some neurons behave differently. Thus, astrocytes convert hyper synchronization state to chimera (Third row (b)). Here, the value of ***N*** = 1000 and ***R*** is 350.

The introducing astrocyte into the network significantly alters neuronal behaviour, leading to an increased desynchronization among neurons. This change is further substantiated by the mean ISI analyses presented in Fig. 8(b) and 9(b), which illustrate the impact of astrocytes on neuronal firing patterns. In the second rows of Figures 8 and 9, the network exhibits chimera states at σ = 0.052 and σ = 0.073. In this scenario, the presence of astrocytes contributes to greater desynchronization among neurons and results in a broader variation in their ISI values. This variation indicates that astrocytes play a critical role in modulating the synchronization dynamics of the network. Finally, in the last rows of Figures 8 and 9, we observe that astrocytes facilitate the transition from hyper-synchronized states (at σ = 0.122 and σ = 0.15) to chimera states by creating tripartite synapses. This mechanism leads to forming of groups of neurons exhibiting equal ISI, thereby confirming the regulatory effect of astrocytes on neuronal dynamics. The findings highlight the complex interplay between astrocytes and neurons, suggesting that astrocytes not only support neuronal function but also actively influence the timing and variability of neuronal firing, which can have significant implications for understanding brain function and potential therapeutic approaches for neurological disorders.

In Figures 10 and 11, the (*x*_*n*_, *y*_*n*_)-plane, or phase plane, of 1000 neurons for varying coupling strengths is illustrated, with different colours. The figures depict two scenarios: one without the presence of astrocytes (panel (a)) and one with astrocytes (panel (b)). We considered three distinct cases of coupling strengths for R = 250 (σ = 0.052, 0.122, and 1) and for R = 350 (σ = 0.073, 0.15, and 1). The analysis reveals that while the presence of astrocytes does not significantly alter the overall dynamics of neuronal behaviour, it does influence the phase plane of specific neurons, causing them to become more spatially separated from one another. In the case of lower coupling strengths (σ = 0.052 in Fig.10 and σ = 0.073 in Fig.11), the effect of astrocytes on the phase plane of neurons is pronounced, leading to substantial changes in the spatial arrangement of neuronal states. Conversely, at higher coupling strengths (σ = 1 in both Figures 10 and 11), the effect of astrocytes on the phase plane becomes less pronounced, indicating that their influence diminishes as the coupling strength increases. This observation suggests that the role of astrocytes in modulating neuronal dynamics is more critical at lower coupling strengths, where their effects can lead to substantial variations in neuronal behaviour. Conversely, at higher coupling strengths, the interactions among neurons may overshadow the regulatory influence of astrocytes, resulting in a more uniform phase plane. The separation of neurons in the phase plane may reflect an increased diversity in firing patterns, potentially enhancing the network’s capacity for information processing. Understanding these dynamics is necessary for elucidating the complex interplay between astrocytes and neurons, which may have significant implications for neural network function. Figure 12 (a) and (b) illustrates how astrocytes can facilitate synaptic plasticity, particularly STDP by forming coherent and coherent regions through chimera states. Coherent regions are associated with Long Term Potentiation (LTP), meaning in this region synaptic strength increased, communication between neurons enhanced, and finally learning and memory will occur. Incoherent regions are related to Long Term Depression (LTD), meaning synaptic strength decreased, weaker communication between neurons exists, and finally leads to selectively weaken specific synapses to make constructive use of LTP. Consequently, LTP strengthens connections between neurons, leading to increased coherence in their activity (coherent region). On the other hand, LTD can weaken connections, resulting in decreased coherence and more desynchronized activity (incoherent region).

**Figure 10:**
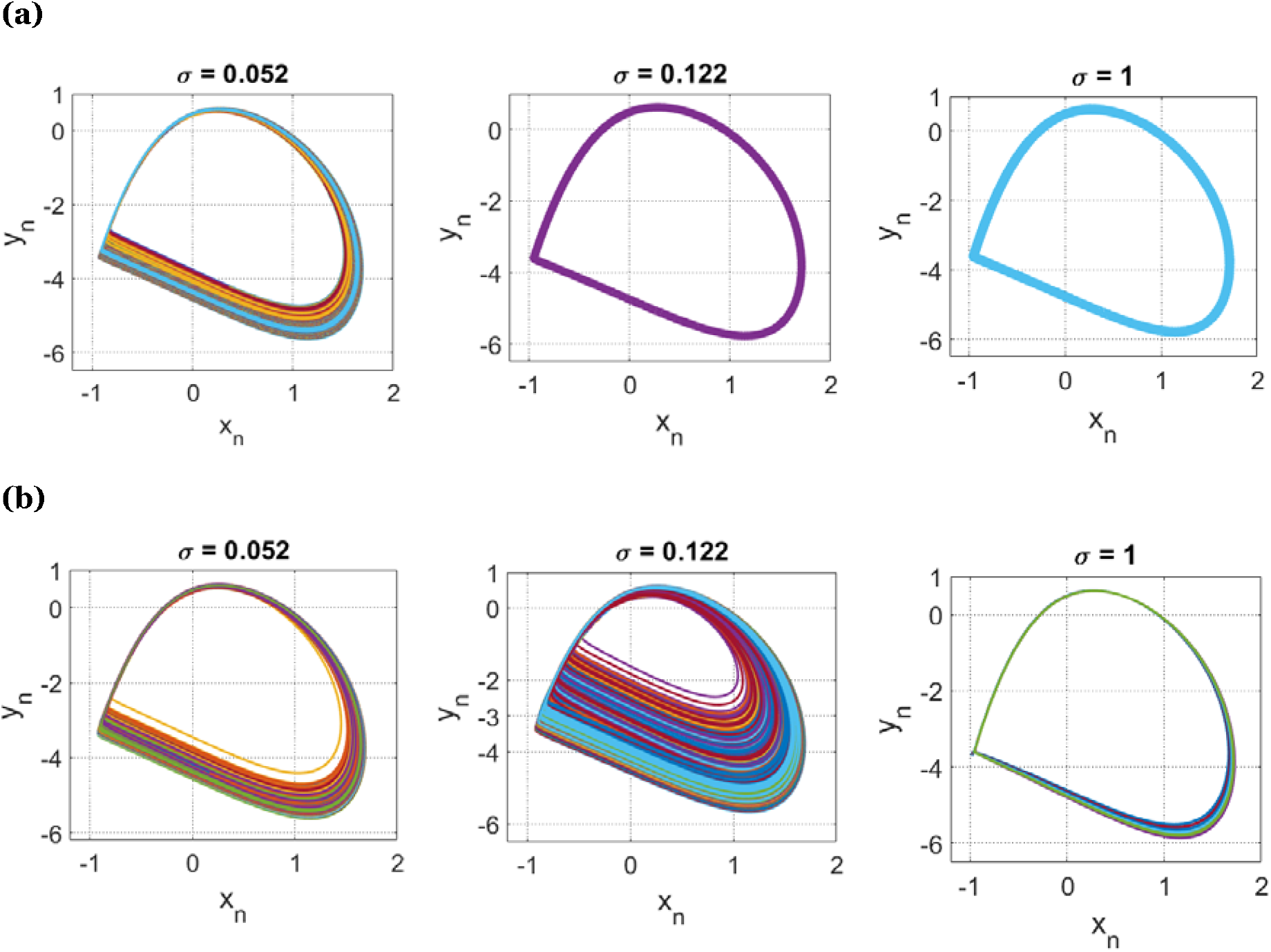
The 2D phase portraits of the neurons with and without astrocytes with increasing coupling strength (), showing different behaviours of the network during 2760mSec - 2840mSec. **(a)** Phase-plane for each neuron with different colour by considering = o.o52, 0.122, and 1 without the presence of astrocytes. **(b)** Phase-plane for each neuron by considering the same coupling strength when astrocytes included. Here ***N*** = 1000 is the number of neurons in the network and ***R***=250. Astrocytes significantly alter the phase plane of neurons when the coupling strength (σ) is set to 0.052. However, at a higher coupling strength (σ = 1), while astrocytes still exert some influence on certain neurons, their impact is considerably less pronounced.

**Figure 11:**
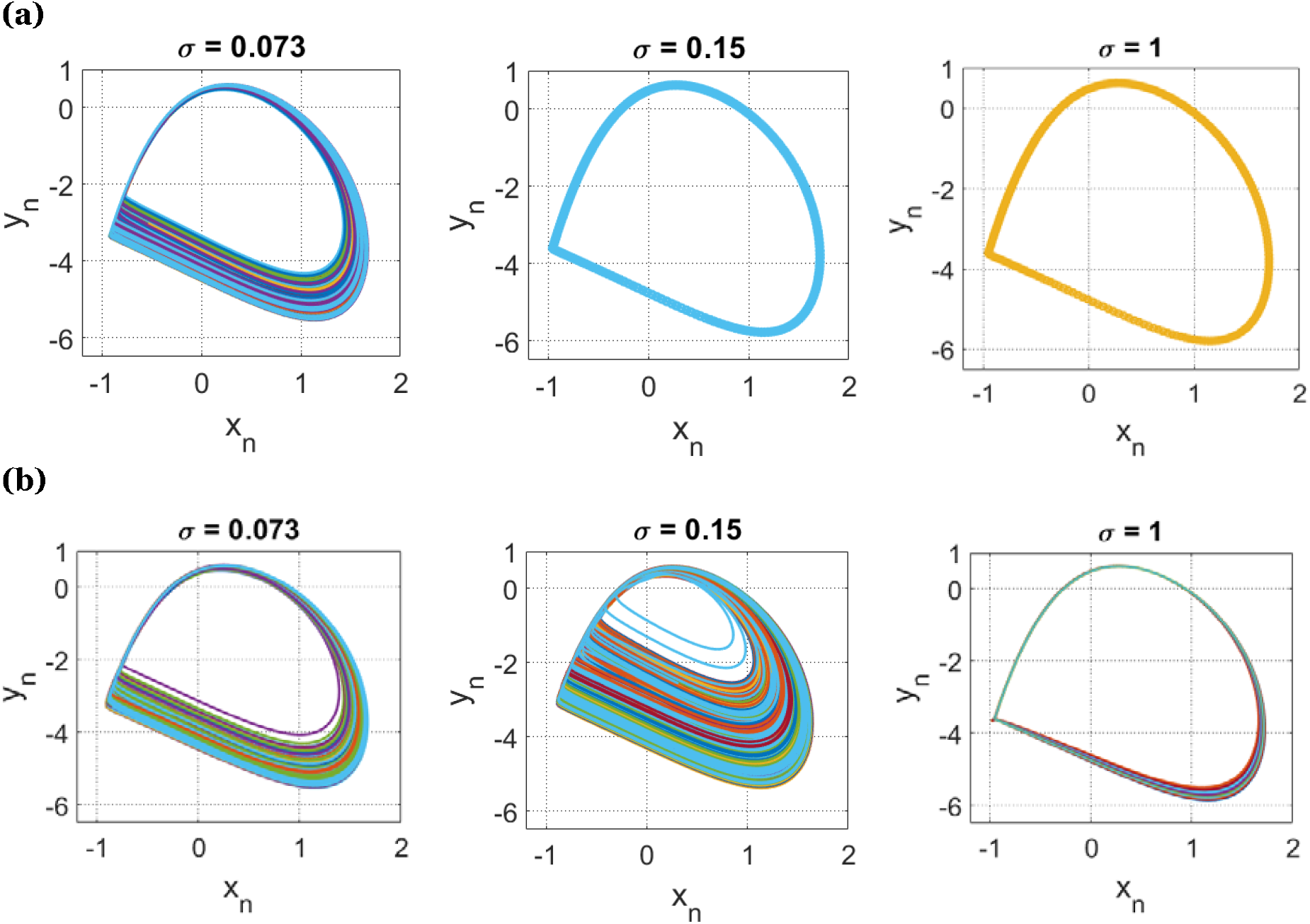
The 2D phase portraits of the neurons with and without astrocytes with increasing coupling strength (), showing different behaviours of the network during 3900mSec - 4000mSec. (a) Phase-plane for each neuron with different colour by considering = o.o73, 0.15, and 1 without the presence of astrocytes. (b) Phase-plane for each neuron by considering the same coupling strength when astrocytes included. Here N = 1000 is the number of neurons in the network and R=350. Astrocytes significantly alter the phase plane of neurons when the coupling strength (σ) is set to 0.073. However, at a higher coupling strength (σ = 1), while astrocytes still exert some influence on certain neurons, their impact is considerably less pronounced.

**Figure 12:**
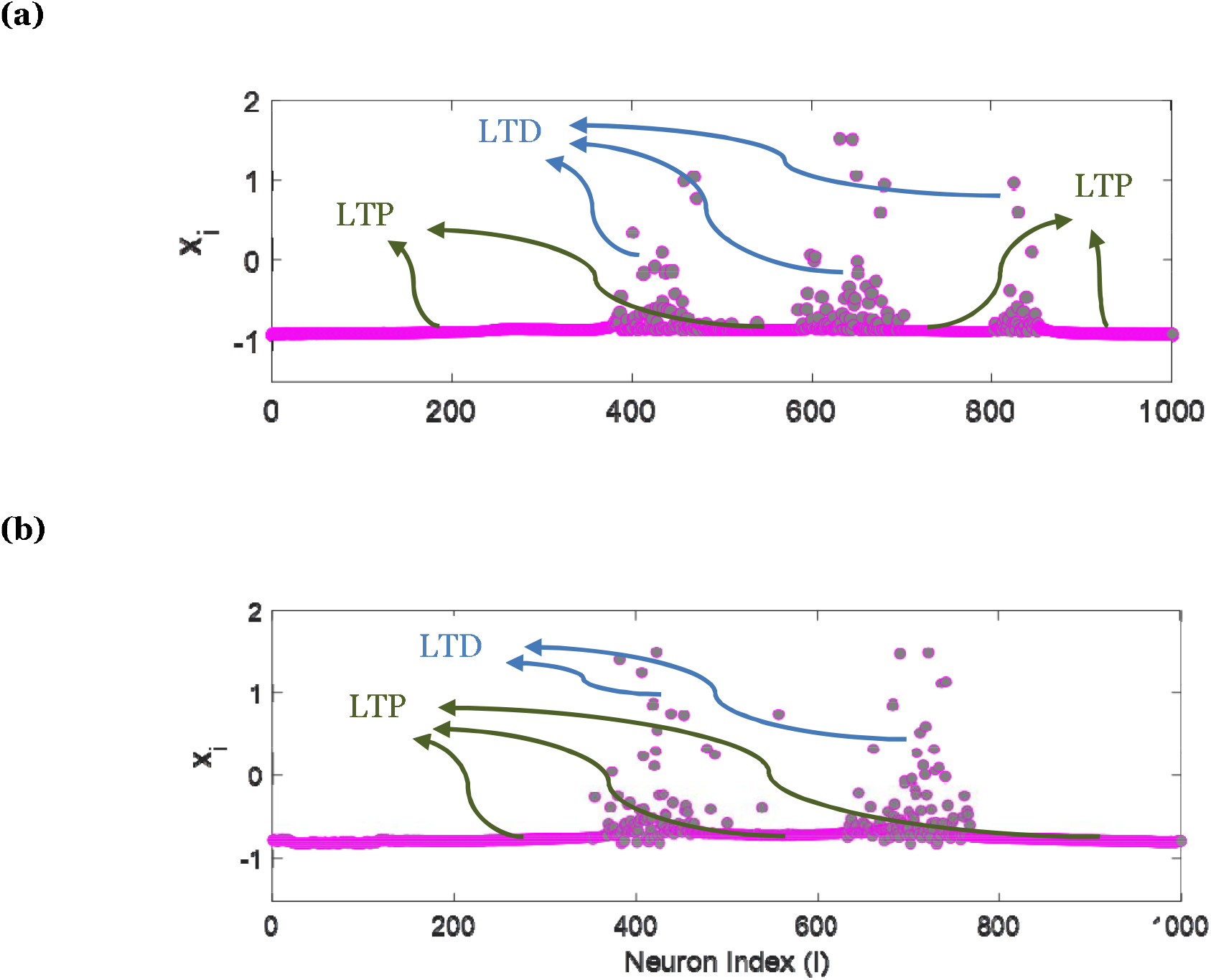
Temporal snapshot of neural and astrocytic activity at 2870 mSec, showcasing the interactions of 1000 neurons and 500 astrocytes, highlighting regions of Long-Term Potentiation (LTP) and Long-Term Depression (LTD), which correspond to coherent and incoherent neural activity regions, respectively. **(a)** with R=250. **(b)** R=350.

## 5. Conclusion

Chimera states represent a fascinating area of research with significant implications for neuroscience. Previous studies focusing on the emergence of chimera states have been conducted on various networks utilizing different neuron models, such as Hodgkin-Huxley, FitzHugh-Nagumo (FHN), the integrate-and-fire model, the Hindmarsh–Rose (HR) model, and the Izhikevich neuron model. However, none of these studies have considered the role of astrocytes in inducing chimera states, despite recent biological findings confirming that astrocyte-neuron crosstalk enhances information processing and learning mechanisms in the brain. By focusing on the interplay between astrocytes and neural synchronization, this research could significantly improve our understanding of brain dynamics and inform future therapeutic strategies. In this paper, we developed a spiking neuronal network that interacts with astrocytes comprising 1000 neurons and 500 astrocytes Under specific model parameters (**λ**_1_ and **λ**_2_) influenced by feedback from astrocytes, the network exhibited robust chimera states. Notably, astrocytes facilitated the transition from hyper-synchronized states to healthy chimera states, which are considered the network’s normal state. The model incorporates astrocyte activation, based on the integrative level of neuronal firing and astrocyte-to-neuron feedback, which relies on the release of the glutamate to promote group spiking of neurons inside the astrocyte’s domain. In this study, we examined and elucidated the underlying mechanisms by which astrocytes facilitate the conversion of hyper-synchronization to healthy chimera states. Utilizing time spans, spatiotemporal analysis, distributions of interspike intervals (ISI), and phase plane diagrams of 2D H-R neurons, we confirmed our hypothesis regarding the critical role of astrocytes in the emergence of chimera states. Our findings indicate that astrocytes significantly influence learning mechanisms by creating regions of coherence and incoherence by formation of chimera states within the neuronal network. This phenomenon is closely related to long-term potentiation (LTP) and long-term depression (LTD) in spike-timing-dependent plasticity (STDP), highlighting the importance of astrocytic activity in modulating synaptic dynamics and enhancing cognitive functions [47, 48]. These results may pave the way for innovative therapeutic approaches aimed at restoring normal neural activity patterns, ultimately enhancing patient outcomes in conditions such as epilepsy and Parkinson’s disease. This research not only contributes to the understanding of neural dynamics but also offers a promising avenue for future clinical applications.

## Compliance with Ethical Standards

### Author contribution

F.A and S.N. developed the theoretical formalism, F.A wrote the final version of the manuscript, performed the analytic calculations and the numerical simulations. All authors investigated the results and write the final version of the manuscript.

### Funding

This study was not funded by anywhere.

### Data Availability

Data would be available through corresponding author with reasonable request.

### Ethical approval

This article does not contain any studies with human participants or animals performed by any of the authors.

